# Low-Cost, Rapid Fabrication of Customizable Polyethylene Glycol-Based Cell Culture Devices

**DOI:** 10.64898/2025.12.04.692348

**Authors:** Emily L. Pallack, Maxwell W. Oulundsen, Hannah R. Goldberg, Yan Kolpakov, Aneth J. Fernandez, Noah D. Teaney, Faith E. Y. Moran, Nisha R. Iyer

**Affiliations:** Department of Biomedical Engineering, Tufts University, Medford, MA, USA

**Keywords:** 3D printing, casting, hydrogel device fabrication, human pluripotent stem cells, neurite extension

## Abstract

Biological research groups may face a high barrier to entry when constructing custom 3D cell culture devices to investigate multi-tissue interactions *in vitro*. Standard fabrication methods such as lithography, etching, or molding are expensive and require specialized equipment and expertise. To address this, we developed an accessible approach for producing polyethylene glycol (PEG)-based cell culture devices using stereolithography (SLA) 3D printing with a polydimethylsiloxane (PDMS) intermediate mold. Both the intermediate molding steps and the sterilized final device show low cytotoxicity, the final device swells to predictable dimensions and retains its shape for at least 10 days. We used this approach to develop a human pluripotent stem cell (hPSC)-derived neural spheroid outgrowth model that supports directed neurite extension over 14 days. Together, this method provides a highly customizable, affordable platform for rapid fabrication of PEG-based microphysiological systems (MPS) for diverse tissue engineering applications.

**Impact:** As biomedical labs work to complement animal models with tissue-engineered MPSs, there is a growing need for low-cost, rapid, and iterative fabrication workflows. We developed a pipeline combining 3D printing, a PDMS intermediate mold, and PEG casting, avoiding the need for specialized photolithography. The resulting devices support stable, nutrient-permissive cell culture while allowing control over device dimensions and customizable channel or compartment configurations. We demonstrate its utility with reprogrammed hPSC-derived neurons, which remain challenging to support sustained neurite outgrowth in engineered models. This workflow expands access to cell culture device fabrication for MPSs across a broader range of biological laboratories.

## Introduction

With growing impetus to move away from animal models and toward *in vitro* models that capture human-specific biology, there is a pressing need for more tissue-engineered microphysiological systems (MPSs) capable of modeling diverse tissue types and developmental processes. While commercial MPS devices are increasingly available, these are often application-specific and not customizable (1,2). These challenges are amplified for multi-tissue devices that demand rapid, iterative design for optimization. However, conventional methods for constructing devices, including photolithography, soft lithography, silicon chip etching, and injection molding require highly specialized clean room facilities and equipment (3–6). As a result, customized devices remain inaccessible to biological research groups that often lack the expertise, budget, and time to prototype bespoke MPSs.

Stereolithography (SLA) 3D printing has been used to overcome these barriers by enabling rapid, low-cost fabrication of molds for facile casting of PDMS, a material valued for device fabrication for its tunable stiffness, gas permeability, optical transparency, and affordability. However, PDMS molds can be cytotoxic due to residual unpolymerized monomers and photoinitiators present in proprietary resin formulations (7–9). While post-processing treatments have been used to shield contaminants from cell-facing materials, the results are inconsistent across resins or require additional expensive and highly specialized equipment to ensure cytocompatibility (8,10,11).

PDMS may also have limitations for certain applications, as it can adsorb small hydrophobic molecules, reducing the effectiveness of biochemical cues in cell media (12–14). Polyethylene glycol (PEG) provides a tunable, affordable alternative to PDMS and is widely used as a hydrogel in cell culture (15,16). UV-curable polyethylene glycol dimethacrylate (PEGDMA) forms cytocompatible, hydrophilic devices that support nutrient diffusion (15,16). PEGDMA devices have been patterned by projection photolithography using digital micromirror devices (DMD) and UV masks (16), or by casting PDMS molds developed by conventional soft lithography (17,18). However, SLA 3D printing has not yet been applied to fabricate PEG-based devices.

Here we propose a novel three-stage molding process using SLA 3D printing and PDMS mold fabrication to produce PEG-based cell culture devices. This approach integrates resin post-processing steps that prevent PDMS curing inhibition, establishes reusable PDMS molds, and relies on equipment accessible to most biological laboratories. We validate the fidelity and reproducibility of feature transfer across the fabrication steps, confirm the cytocompatibility of the materials, and demonstrate the utility of the platform using hPSC-derived neural spheroids for multimillimeter-range neurite outgrowth experiments. Together, these results provide a practical and accessible fabrication strategy for producing PEG-based cell culture device suitable for diverse MPS applications.

## Methods

### Device Fabrication and Printing

Devices were designed on AutoDesk Fusion 360 or TinkerCAD, prepared for printing using Lychee Slicer, and printed using High Resolution Phrozen High-Resolution Aqua 4K Resin (Phrozen, Hsinchu City, Taiwan) on the ELEGOO Mars 3 4K Resin Liquid Crystal Display (LCD) 3D Printer (ELEGOO, Shenzhen, Guangdong, China) (Figure 1A). Phrozen 4K resin was selected for mold fabrication because prior work by Hagemann et al. demonstrated that, after standard post-processing, this resin does not inhibit PDMS curing and therefore requires no additional coating or surface-treatment (8). The ELEGOO Mars 3 printer was selected for its affordability (<$200), sufficient build volume (143 X 89.6 X 175 mm^3^), and resolution (35 µm XY and 20 µm Z layer height).

**Figure 1:**
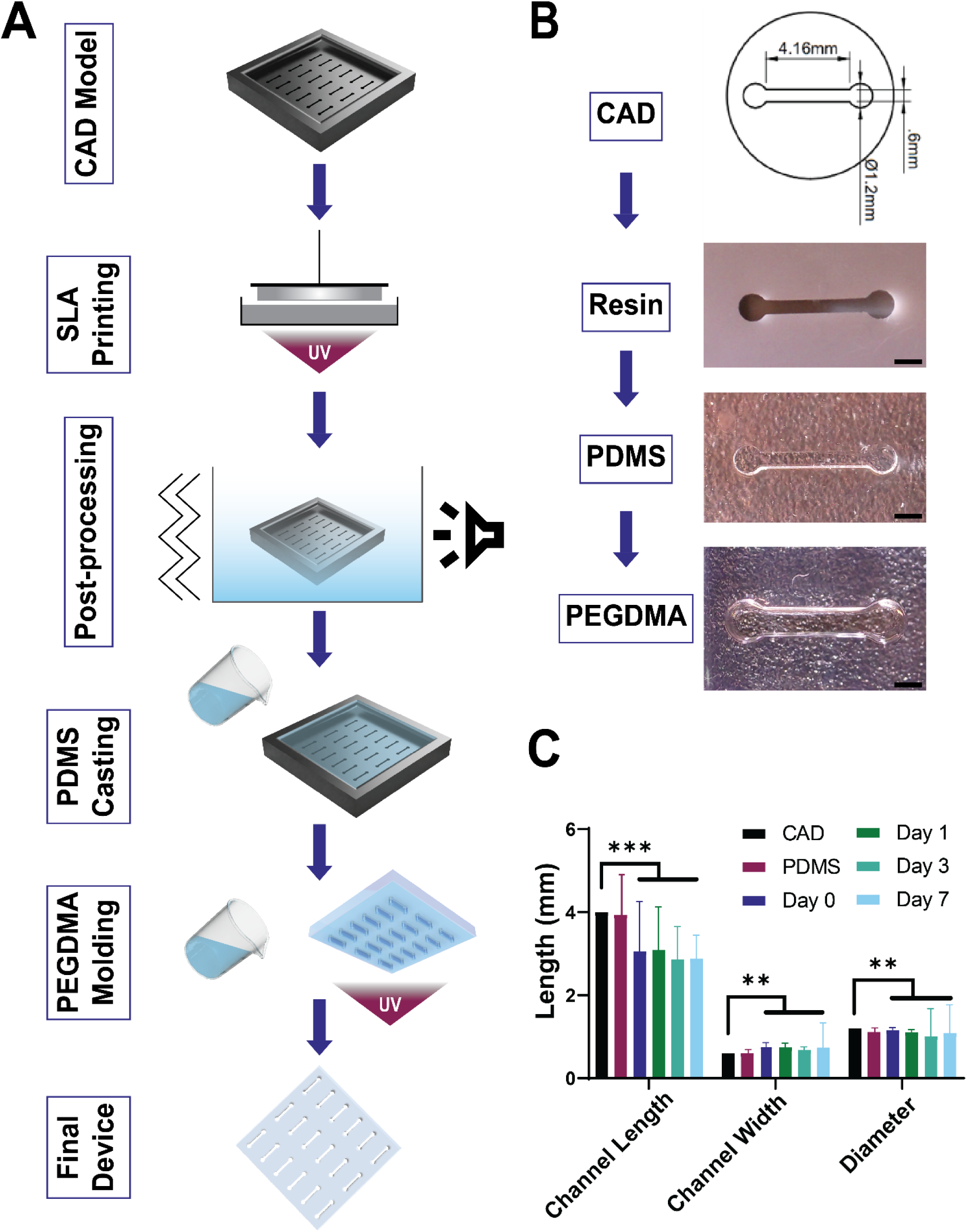
Rapid production of versatile PEG-based cell culture devices. (A) Schematic overview of the protocol to generate final PEGDMA device (B) Representative images of CAD technical drawing, SLA-printed resin mold, PDMS intermediate, final PEGDMA device. Scale = 1mm (C) Comparison of predicted CAD model dimensions to PDMS intermediate and PEGDMA swelling profile over 7 days. *** = p < 0.001, ** = p < 0.01. One way ANOVA with post-hoc Dunnett’s test compared to CAD.

### Resin Print Post-processing

Resin prints were post-processed by curing with 405-410nm UV light for 60 min at 60 °C, washing twice with fresh isopropanol for 20 min and sonicating in isopropanol for 5 minutes, before curing again at 60 °C overnight. The next day, Dow SYLGARD™ 184 Silicone (PDMS, Dow Chemical Company, Midland, MI) was mixed in a 10:1 ratio of base to curing agent by weight. This mixture was degassed under vacuum for 10 minutes, then PDMS was poured into molds and degassed for an additional 10 min. Molds were incubated at 60°C for 4h to allow PDMS to cure, after which the inverse PDMS casts were promptly demolded. After fabrication, PDMS casts were immediately used for PEGDMA casting or stored in a cool, dry place for later use.

### PEG Device Casting and Sterilization

To cast PEGDMA devices, a solution of 10% (w/v) polyethylene glycol dimethacrylate M.W. 1000 (PEGDMA; Polysciences Inc., Warrington, PA, USA) and 1.1 mM lithium phenyl-2,4,6-trimethylbenzoylphosphinate (LAP; Advanced BioMatrix, Carlsbad, CA, USA) was prepared in Dulbecco’s phosphate buffered saline (DPBS, pH 7.4; GIBCO, Waltham, MA, USA). This solution was continually mixed using a magnetic stirrer for 1 hour before immediate use or storage in the dark at 4°C. PDMS molds were briefly wiped with ethanol and placed feature-side down onto Transwell polycarbonate membrane inserts (0.4 μm pores). PEGDMA solution was then added to fill the negative space beneath the mold. The Transwell assembly was placed into a well plate, and PEGDMA was crosslinked from below using a 405–410 nm UV lamp (60 W luminous efficacy; 6 W power; 90–250 V) for 10 minutes. After curing, the PDMS molds were removed to reveal the PEG-based device. Before cell culture, PEG-based devices were swelled in UltraPure™ DNase/RNase-Free Distilled Water (ThermoFisher, Waltham, MA, USA) with 1% antibiotic-antimycotic (ThermoFisher) solution and placed under germicidal UV light (254nm) for 1hr. Final sterilization was performed by removal of antibiotic/antimycin solution, a wash with sterile Ultrapure water, and addition of appropriate media with 1% antibiotic-antimycotic.

### hiPSC Maintenance

We used a polyclonal pool of KOLF2.1J human induced pluripotent stem cells (hiPSCs; XY; Jackson Laboratory; passage10-30) that were edited via the piggyBac transposon system to express a doxycycline-inducible NGN2 cassette for direct reprogramming neurons as previously described (KOLF2.1J:pB-TO-NGN2) (19,20). For downstream applications, hiPSCs were washed with 1X DPBS, dissociated to single cells using Accutase (Corning, Corning, NY, USA) for 6 min at 37°C and seeded as needed.

hiPSCs were maintained at 37°C in 5% CO2 in Essential 8 (E8, ThermoFisher, Waltham, MA, USA) medium on Cultrex reduced growth factor basement membrane extract (R&D Systems) with routine passaging to maintain pluripotency. hiPSCs were passaged at 70-80% confluency. Briefly, cells were first washed with 1X DPBS and incubated at 37°C with Versene (Invitrogen, Carlsbad, CA, USA) for 6 min. Versene was removed, cells were resuspended with E8 and 50nM Chroman 1 and distributed to new Cultrex-coated 6-well plates at 1:12. E8 media was replaced daily.

### PiggyBac Transfection for Live Imaging

For fluorescent live imaging, KOLF2.1J:pB-TO-NGN2 hiPSCs were transfected with pB-CAG-eGFP (gift from Joseph Loturco (21); Addgene plasmid #40973) or pB-CAG-tdTomato (gift from Mario Capecchi; Addgene plasmid #133569). At ∼50% confluency, culture media was replaced with OptiMEM (ThermoFisher) containing a 1:3 mixture of Super PiggyBac Transposase (System Biosciences, Palo Alto, CA, USA) and the fluorescence plasmid, 5 µl Lipofectamine (ThermoFisher), and CEPT cocktail (50 nM Chroman-1(Tocris), 5 µM Emricasan (Selleck), and 1× Polyamine (Tocris)) as previously described (22). Cells were incubated overnight. The following day, cultures were checked for fluorescence and then returned to E8 media for expansion and subsequent differentiation.

### hPSC Differentiation

To generate cortical neurons, eGFP+ or tdTomato+ KOLF2.1J:pB-TO-NGN2 hiPSCs were first washed with 1X DPBS, dissociated to single cells using Accutase for 6 min at 37°C and seeded at 100k cells/cm^2^ in Essential 6 media (E6, Thermo Fisher, Waltham, MA, USA) containing 2ug/mL doxycycline (DOX, Millipore Sigma, San Diego, CA, USA), 10 µM DAPT (ThermoFisher, Waltham, MA, USA) and CEPT cocktail as previously described(23). The following day, media was changed to E6 with 2ug/mL DOX and 10 µM DAPT kept on for 6 days with daily media changes when neuron morphology was observable by fluorescence microscopy (**Figure 3B**).

### Cytotoxicity Assay

To differentiate cortical neurons for cytotoxicity testing, hiPSCs were plated at 100k/cm^2^ in 24 well plates, followed by DOX induction as described (2ug/mL DOX and 10 µM DAPT for 6 days). Neurons were allowed to recover in 1mL maturation media (E6 with 1X N2, 1X B27, DAPT, and 1% antibiotic–antimycotic) for 2 days with media change daily. PDMS or PEG squares (2 cm²) were floated in culture media for 24 hours, cytotoxicity was assessed using the Molecular Probes LIVE/DEAD Viability/Cytotoxicity kit according to the manufacturer’s instructions. Briefly, neurons were first gently washed with sterile DPBS. EthD-1 stock (2 mM) was diluted to 4 µM in DPBS and vortexed, followed by addition of calcein AM (4 mM stock) to a final concentration of 2 µM and vortexing again. A total of 250 µL of staining solution was added per well and incubated at room temperature for 30 minutes. Cells were imaged immediately on an EVOS microscope using GFP and RFP filter cubes (three 10× fields per well). Viability was quantified in ImageJ by manually thresholding green (live) and red (dead) signals, and percent viability was calculated as green area divided by the sum of green and red areas.

### Spheroid Formation and Device Seeding

To seed devices, neural spheroids were formed via the hanging drop method as previously described (24). Briefly, 10 µL of singularized neurons (approximately 25,000 cells/spheroid) in E6 media containing 1% antibiotic–antimycotic and 50 nM Chroman 1 were placed on the inner lid of a 96-well plate. Lids were inverted over wells prefilled with E6 to prevent dehydration and incubated at 37°C, 5% CO₂ for 3 days. Using a digital microscope for guidance, spheroids were lightly aspirated with a P200 pipette and placed into device compartments. Excess media was removed with a Kimwipe, and approximately 20 µL of 1:1 Cultrex:E6 was added to backfill compartments, then incubated at 37°C for 15 minutes for gelation. Finally, 1 mL of maturation media (E6 with 1X N2, 1X B27, DAPT, and 1% antibiotic–antimycotic) was added to the lower compartment of the Transwell system and changed daily for 14 days.

### Immunostaining

PEG-based devices were removed from Transwells, separated with a razor, and placed individually into 24-well plates. Devices were fixed in 4% paraformaldehyde for 30 min at room temperature, then washed three times with 1 mL Tris-buffered saline (TBS, ThermoFisher). Samples were blocked and permeabilized in TBS containing 5% donkey serum and 0.3% Triton X-100 for 1 hour at room temperature. Primary antibody against TUJ1 (Biolegend, Rabbit, 1:1000) diluted in TBS with 5% donkey serum and 0.3% Triton X-100, was applied overnight at 4°C. Devices were washed three times with TBS + 0.3% Triton X-100 for 10 min each. Alexa Fluor secondary antibodies (1:500), sterile-filtered through a 0.22 µm filter, were applied for 1 hour at room temperature, followed by two 15-minute washes with TBS. DAPI (1:2000 in TBS) was added for 10 min at room temperature, then washed once for 15 min with TBS. Devices were imaged at 4X on a Molecular Devices ImageXpress Micro Confocal High-Content Imaging System.

### Statistics

Statistical analyses were performed in JMP Student Edition 18.2.2. Graphs were constructed using GraphPad 8.4. Device characteristics (channel width, length, diameter, cytotoxicity) were compared using one-way ANOVA with Dunnett’s post-hoc test against CAD designs or control neuron cultures. All assays were performed with three independent biological replicates. To evaluate changes in neurite extension over time and determine whether growth trajectories differed between groups, we performed a two-way ANOVA with Day (continuous) and Group (eGFP vs TdTomato) as fixed effects and included the Day × Group interaction term to assess differences in growth rate. *** = p < 0.001, ** = p < 0.01

## Results

### Novel Fabrication of PEG-based Devices with Predictable Geometries

We developed a three-stage molding workflow to fabricate PEG-based devices with precise geometries (**Figure 1A,B**). First, we designed the desired geometry in CAD and tiled multiple devices to increase throughput and reduce print time. The model was then sliced and fabricated via SLA 3D printing. After post-processing, PDMS was cast into the resin mold to create an inverse mold, which was subsequently used to generate the final PEG-based device. To assess feature fidelity, we replicated a representative “barbell” geometry—applied in recent innervation(1,25) and neural MPS studies (26)—through the resin, PDMS, and PEGDMA steps (**Figure 1B**), confirming consistent geometry transfer and reproducibility across materials and fabrication runs. Quantitative analysis showed significant changes in channel length, width, and compartment diameter in the final PEG-based device compared to the original CAD model; however, swelling remained consistent and stabilized within 1 day in aqueous conditions (**Figure 1C**). Based on these results, we anticipate the need to design channel lengths ∼20% longer in CAD models to account for PEGDMA swelling in confined geometries.

### Reusable Molds Enable Affordable Device Production

To determine whether PDMS molds could be produced with different conformations and without repeating the full resin-to-PDMS-to-PEG workflow, we designed T- and L-shaped channel geometries for multiple castings. (**Figure 2**). PDMS cast into resin molds produced first-generation PEG-based devices, which were filled with Trypan Blue to confirm geometric fidelity (**Figure 2**, CAD and 1st Cast). Using the same PDMS molds, second-generation PEG devices were cast with no visible distortion or loss of resolution (**Figure 2**, 2nd Cast), demonstrating that SLA-printed molds and PDMS intermediates can be reused for multiple cycles.

**Figure 2:**
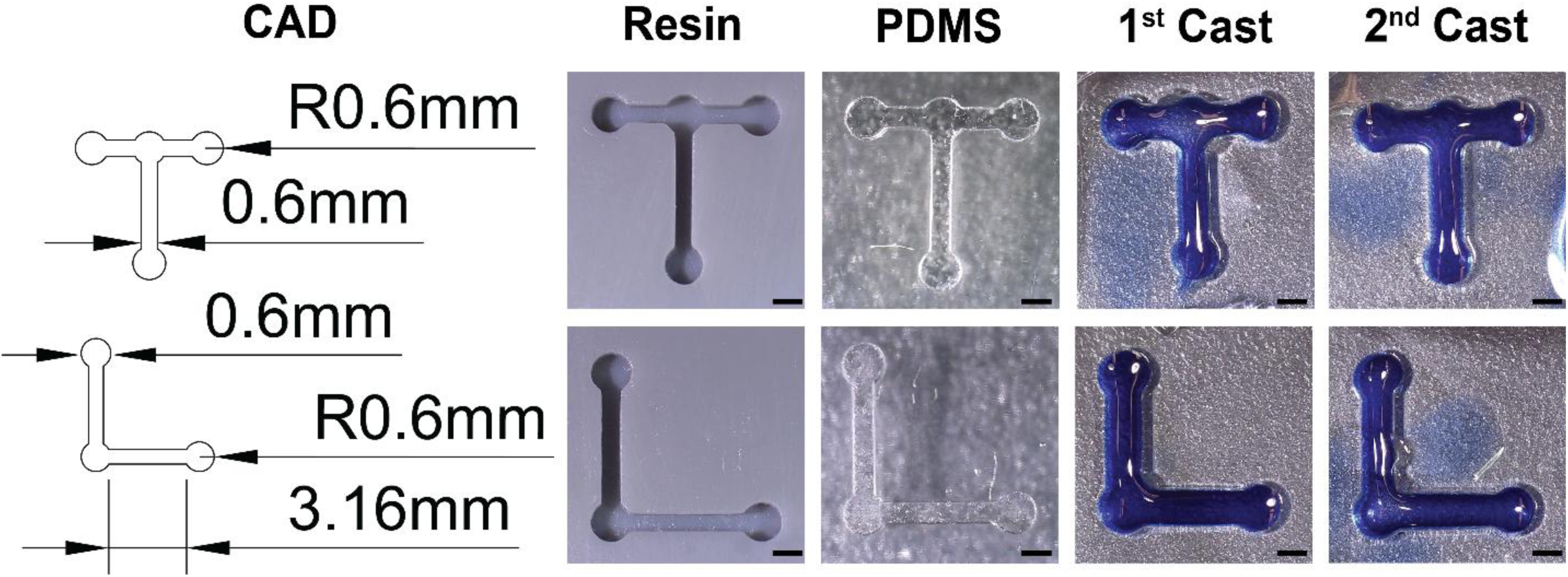
CAD schematic of designed custom configurations. Resin print in Phrozen Aqua Grey 4K and Sylgard 184 PDMS. PEG devices from first and second cast visualized with AlamarBlue. Scale = 1mm

### Intermediate Mold and Final Devices Support Cell Viability

Prior reports suggest resin-derived acrylate leachates can reduce cell survival (27), so we directly tested hPSC-derived neuron viability in cultures exposed to resin-cast PDMS or PEGDMA for 24 hours (**Figure 3A)**. Live/dead staining showed dense fields of viable cells across control, PDMS-, and PEGDMA-conditioned groups, with few dead cells in any condition (**Figure 3C**). Quantification confirmed high viability (>90%) and no statistically significant differences among groups (**Figure 3D**). The high viability observed here likely reflects thorough resin post-processing during PDMS curing, which minimizes residual cytotoxic components.

**Figure 3.**
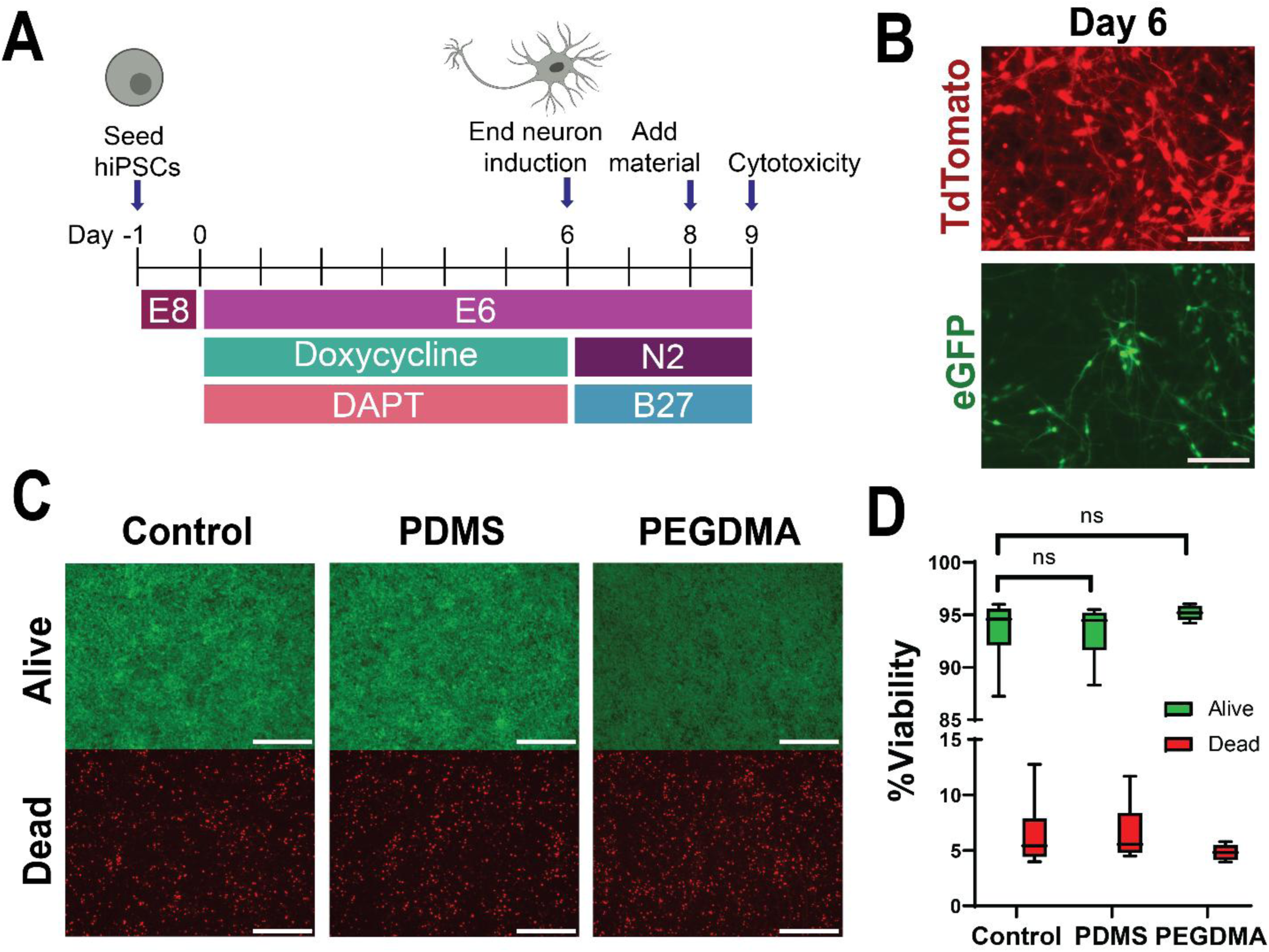
Final and intermediate device materials are not cytotoxic. (A) Experimental schematic for cytotoxicity study (B) Representative hiPSC-derived neuron morphology at day 6 of induction. Scale = 150 µm (C) Representative images of iPSC direct reprogrammed neurons labeled with calcein (live, green) and propidium iodine (dead, red) after 24 hours of soluble exposure to material. Scale = 300 µm. (D) Quantification of iPSC direct reprogrammed neuron viability at 24 hours (percent area green or red versus additive green and read area). 3 independent wells imaged in 3 different locations per condition. A one-way ANOVA with post-hoc Dunnett’s test was used to compare PDMS and PEGDMA to neuron-only control (control vs PDMS p= 0.9969, control vs PEGDMA p=0.1865)

### Two-Compartment Device Promotes Controlled Neurite Extension

Applying our fabrication workflow, we modeled hPSC-derived neurosphere co-culture and evaluated neurite extension over 14 days. Fluorescent tdTomato- and eGFP-expressing cortical neurons were differentiated and aggregated (**Figure 4A**), then spheroids were placed in adjacent device reservoirs (**Figure 4B**) and stabilized with Cultrex to provide an extracellular matrix conducive to neurite growth (**Figure 4B**). Live imaging confirmed successful loading on Day 1 (**Figure 4C**). By Day 10, dense, aligned neurite bundles extended along the channel, forming continuous axonal bridges with minimal spheroid migration outside the channels (**Figure 4C**). Quantitative analysis showed neurite length increased 2-3 mm over 14 days, aligned within the PEGDMA channel, which remained intact (**Figure 4D**).Two-way ANOVA revealed a significant effect of time (p < 0.0001) but no main effect of group (eGFP vs TdTomato) (p = 0.1388), with a significant Day × Group interaction (p = 0.0081) indicating faster neurite extension in the eGFP+ population. Slight neurite die-back was observed at Day 14, likely due to the absence of exogenous growth factors in the media. Finally, to assess the feasibility of whole-mount staining, devices were immunolabeled for TUJ1 and imaged with confocal microscopy, confirming neuronal identity throughout the device (**Figure 4D**). This demonstrates that our PEGDMA platform supports robust, spatially controlled neurite extension in multiple compartments, enabling both live imaging and immunolabeling of fixed samples for versatile analysis of neuronal connectivity *in vitro*.

**Figure 4.**
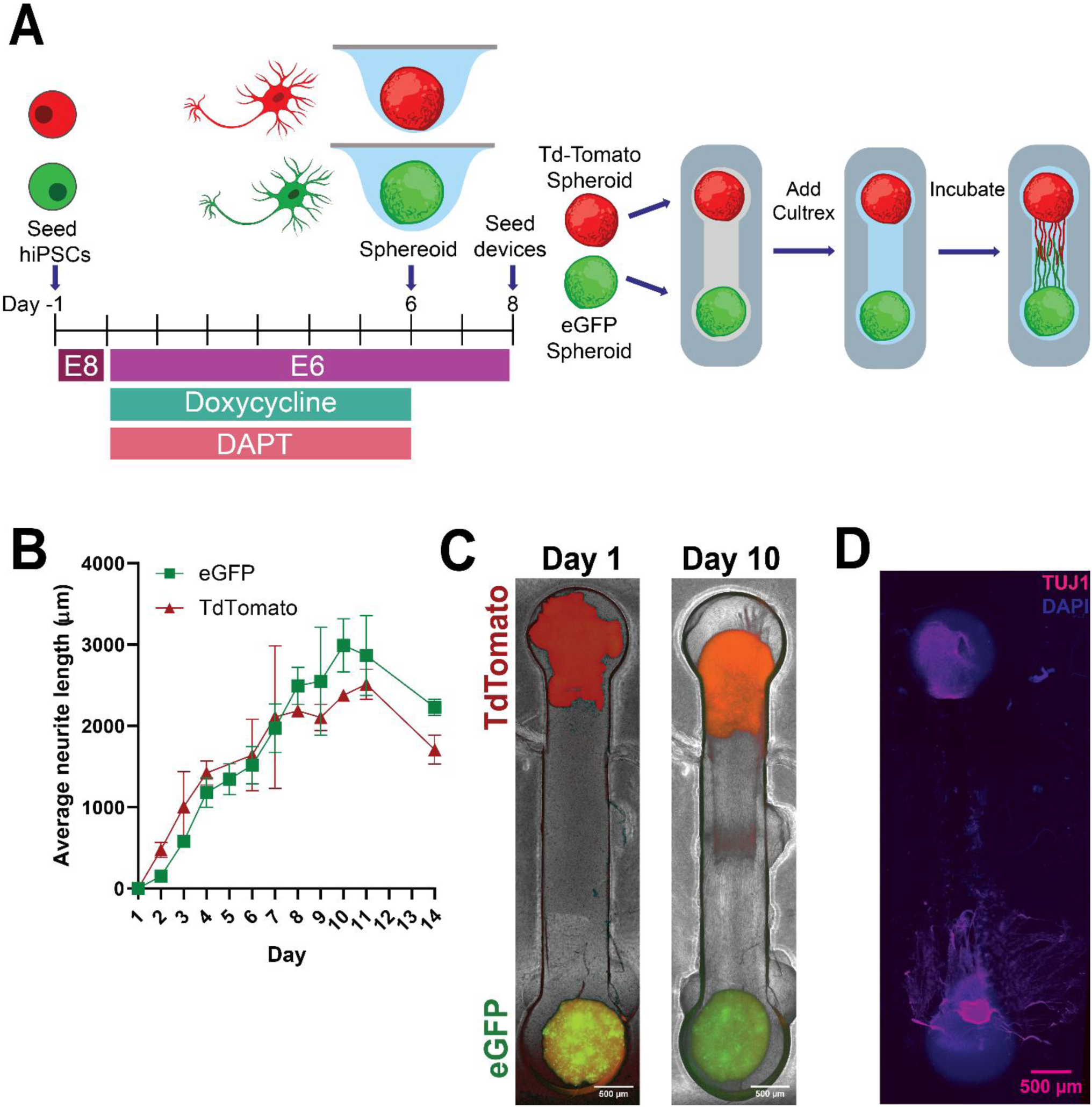
PEG-based device with Cultrex infill supports robust neurite outgrowth. (a) Differentiation timeline of neural spheroids and experimental set up (b) Quantification of neurite length in PEG-based cell culture device measured over 14 days. Error bars are mean with +/- SEM. (c) Live fluorescence imaging of neural spheroid device constructs at day 1 and day 10 of confined neurite outgrowth through Cultrex. (d) Whole mount staining of TUJ1+ neurite extensions at day 14.

## Discussion

We present a low-cost, rapid method for fabricating PEG-based cell culture devices that can be customized for diverse applications. This protocol builds on existing microfabrication pipelines by integrating 3D printed resin molds, PDMS intermediates, and PEGDMA casting into a single, flexible workflow (8,16,28). Cost analysis indicates that equipment, software, and consumables are affordable (**Table 1**), with a single 4 mm barbell device costing ∼$0.22—further reduced by PDMS mold reuse and CAD tiling to produce multiple molds per print. The workflow is rapid: SLA prints take ∼20 minutes, the only long step is a single overnight cure, and mold reuse greatly shortens fabrication time and turnaround for additional devices.

**Table 1.**
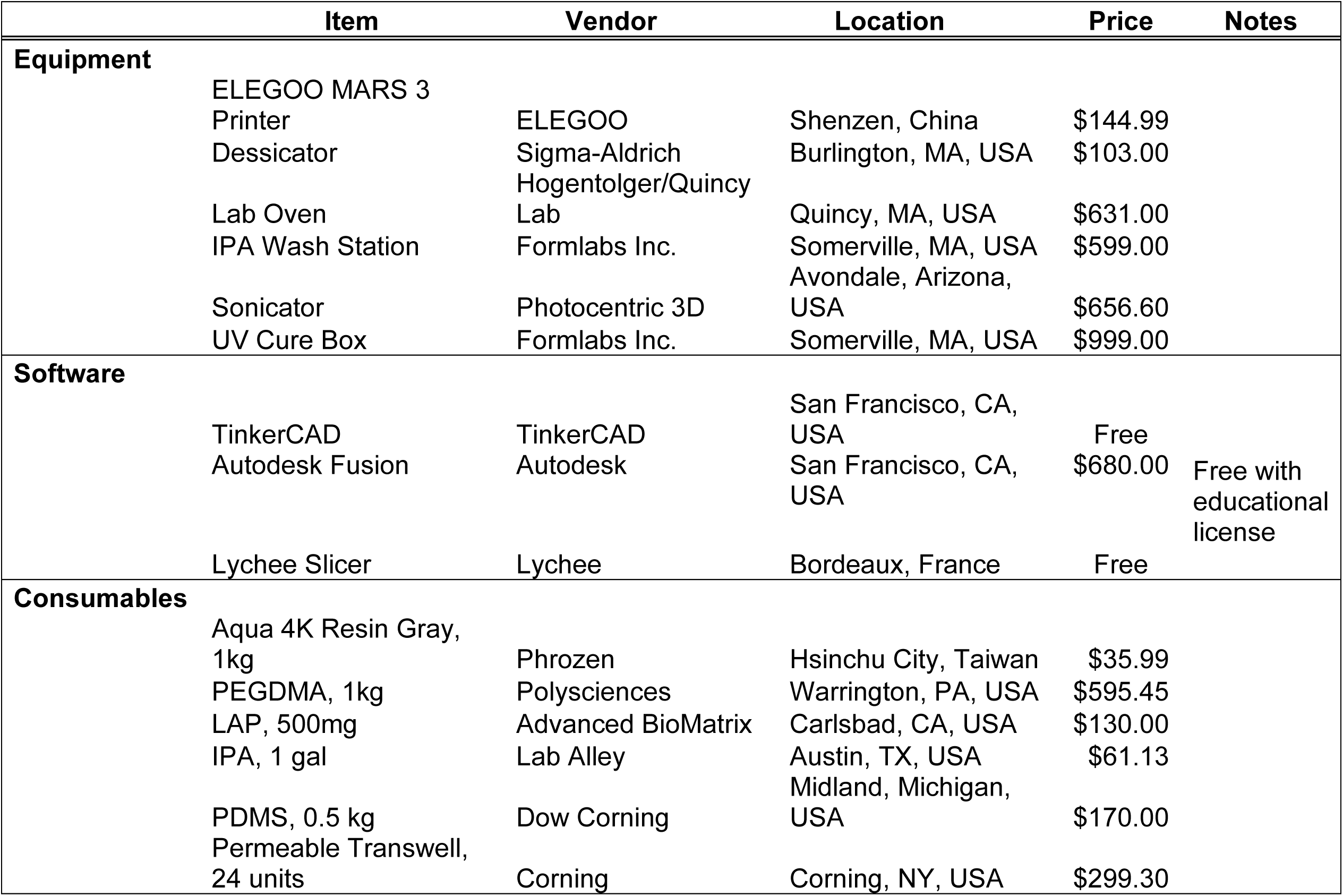
Categorization of standard prices for equipment and consumable items for implementing workflow.

A major caveat to this approach is that device design is limited by resin mold resolution. The ELEGOO Mars 3 used here offers 35 µm XY resolution, sufficient for millimeter-scale channels but not finer microfeatures. Higher-resolution options, such as 8K LCD printers (∼20 µm XY) or Digital Light Processing systems (<10 µm pixel size), allow smaller structures but often at higher cost and reduced build volume. Newer models, like the ELEGOO Mars 5 Ultra, improve both resolution (∼18 µm XY) and build area (153 × 77 × 165 mm). Two-photon lithography, including high-throughput platforms like the UpNano NanoOne, can achieve sub-micron resolution but requires six-figure investments and specialized training, putting it beyond the reach of typical academic or maker-space groups (29) At very small feature scales, demolding becomes challenging: success depends on the ratio of feature depth to overlying PDMS thickness. While smaller features are likely achievable, we did not optimize this ratio, and tall, narrow features surrounded by thicker PDMS can resist release. Researchers should choose a workflow that balances spatial resolution, accessibility, and cost.

Another important consideration is the hydrogel backfill, which can influence both long-term culture stability and overall cost. While Transwells are one of the more expensive components (**Table 1**), they help maintain device integrity during demolding, as noted by others (28), because the Cultrex-in-PEGDMA backfill is not physically entangled and can detach. This limitation could be mitigated by incorporating an inner hydrogel compatible with PEGDMA, such as functionalized PEG variants commonly used in tissue engineering (30) Transwells also support neuronal health and enable long-term culture that may not be feasible on standard tissue-culture plastic or glass, introducing another tradeoff between device design, culture longevity, and experimental use case.

While achievable resolution and material choices may preclude printing of microscale features, such as those in classic Campenot chambers (31), the millimeter-scale dimensions of our PEG-based devices still enable meaningful spatial organization and functional compartmentalization of neuronal cultures. This may benefit 3D organoid and spheroid co-cultures by directing neurite projections across defined microchannels while keeping spheroids spatially anchored, avoiding the manual alignment required for assembloids (32). Our platform provides a structured, customizable architecture for multiple spheroids or organoids in defined compartments. This enables reproducible modeling of spatiotemporal neural development, multi-region communication, and disease-relevant circuitry using distinct cell populations. Here we use hPSC-derived cortical neurons, but similar devices have been applied for rodent (1) and hPSC-derived peripheral sensory neurons (25), primary cortical neurons (33), and astrocytes (24). Beyond the nervous system, the platform could be adapted for other tissue types, offering broad experimental flexibility. Because of the casting-based workflow, additional casting steps could be incorporated to introduce functionalized PEG or other synthetic/ECM hydrogels, enabling modeling of extracellular matrix contributions to development or disease. Finally, by combining consumer-grade SLA printing with PDMS, this workflow is accessible to a wide range of biological laboratories and teaching settings, from high school to undergraduate courses, offering a scalable, practical tool for device fabrication.

## Acknowledgements

The authors would like to thank Angela Lai and Charlotte Jacobus for their continuous feedback and support. We would also like to thank Laura Place for her assistance in equipment access, Edward Gordon for sharing his expertise in 3D printing, and Carmen Preda for administrative support.

## Author Contributions

E.L.P. – Conceptualization, Investigation, Visualization, Software, Writing - original draft, Formal analysis, Project administration; M.W.O – Formal analysis, Investigation, Methodology, Writing – original draft; H.R.G – Formal analysis, Validation, Writing – original draft; Y.K. – Formal analysis, Software, Validation; N.D.T., A.J.F., & F.E.Y.M. – Formal analysis, Investigation, Methodology; N.I. – Conceptualization, Writing – original draft, Writing – reviewing and editing, Supervision, Project administration, Funding acquisition

## Conflict of Interest

The authors declare no competing interests.

## Funding Statement

This work was supported by the Tufts University Undergraduate Research Fund (M.W.O., N.D.T., A.J.F., and F.E.Y.M.), NSF NSFGRFP2023348129 (E.L.P.), Tufts Department of Biomedical Engineering Start-up (N.R.I.), and NIH DP2NS140734-01 (N.R.I.)

## Notes

### Competing Interest Statement

The authors have declared no competing interest.

